# Decision revisions modulated by synaptic inhibition in the olfactory bulb facilitate learning

**DOI:** 10.64898/2025.12.23.696244

**Authors:** Sanyukta Pandey, Arpan Kumar Nayak, Anindya S. Bhattacharjee, Krish Pandey, Susobhan Das, Nixon M. Abraham

## Abstract

Behavioral responses are preceded by decisions based on perceived sensory evidence. In real life, sensory inputs are often noisy and change incessantly, raising the question of when and how accurate decisions are made. Cognitive flexibility allows us to revise the choices in case of perceptual conflicts, leading to response revisions. This involves switching between available choices or rapidly stopping misconstrued responses. Here, we quantified these erroneous behavioral actions and studied the underlying neural mechanisms. Mice were trained on an olfactory decision-making task wherein they had to distinguish between a rewarded and an unrewarded stimulus by responding with a lick or withholding it. While the animals respond by licking for both rewarded and unrewarded odor stimuli at the initial stages of learning, the responses almost disappear for the unrewarded ones in the learned stage. However, animals tend to initiate lick responses and stop within a few milliseconds for a few unrewarded trials, even when accuracy levels are high. We describe this phenomenon as decision revisions (DR), and observed it in 5-25% of trials among high-performance blocks. These revisions were mostly observed within a few hundred milliseconds of stimulation. We observed a significantly higher number of revision trials for binary odor mixture discriminations compared to monomolecular ones. Further, by enhancing inhibitory synaptic signaling in the olfactory bulb through photoactivation of ChR2-expressing GAD65-positive GABAergic interneurons, we observed faster odor discrimination and fewer revision trials. Thus, our findings confirm decision revisions that are stimulus complexity-dependent and the pre-cortical control over such a complex cognitive activity.

**Highlights of the study:**

- Mice revise context-inappropriate responses in a Go/No-go task.
- Mice correct errors as they learn.
- Decision revisions are shaped by stimulus complexity.
- Optogenetic activation of the olfactory bulb inhibitory interneurons modulates decision revisions.

## Introduction

Response revision prompts animals to alter an erroneously initiated action. In a Go/No-Go perceptual decision-making task, the responses depend on the incoming sensory evidence, internal state, and environmental constraints (1–3). Decision speed may affect the accuracy by lowering the evidence thresholds, causing erroneous initial responses (4, 5). These context-inappropriate responses could persist or terminate promptly depending on the extent of error awareness (4, 6, 7). Revisions are likely when awareness about the misjudged context emerges (4, 6). Initial error responses could stem from incomplete sensory evidence, later countermanded through sensory processing and error monitoring (4, 7). Studies have indicated that the brain continues to process information even after response initiation, allowing corrections during difficult tasks (8, 9). This sensory integration post-action initiation leads to the updating of an ongoing response (9). Earlier studies have shown that changes in initial decisions corrected errors more than they impaired the right choices (6). This suggests that a change of mind is highly likely to occur when responses are incorrect. This is driven by continued evidence accumulation, showing flexibility in human decision-making (6, 7).

Change of mind enables response revisions. This is a complex cognitive phenomenon with many brain areas working in tandem facilitating different aspects involved. Neuroimaging studies in humans and primates have shown brain regions involved in various aspects of decision-making and change of mind. Orbito-frontal cortex (OFC), anterior cingulate cortex (ACC), and dorsolateral pre-frontal cortex (DLPFC) control attention (10–12). OFC, along with the striatum, processes reward-based decisions via limbic projections (12). Anterior insula integrates information about salience, uncertainty and emotional aspects of the decisions (13). Temporal and parietal cortices track outcomes, particularly error monitoring and feedback processing (14, 15). Ventro-medial pre-frontal cortex (vmPFC) and hippocampus encode reward values and memory (16, 17). VTA dopaminergic neurons code reward prediction error for cost-benefit decisions (18). Basal ganglia aid decision making by integrating different information and transmitting to the thalamus which further relays it to the cortices (19). The cerebellum modulates decisions by facilitating communication and coordination between brain regions (11). mPFC activity has been linked with error monitoring and change of mind in human subjects (8). Studies have consistently shown that change of mind is inevitable in case of a noisy stimulus, attributing better temporal encoding of stimulus as a hallmark of change of mind (6, 7). However, how sensory processing mechanisms affect the change of mind remains largely unknown.

Rodent studies have utilized the Go/No-Go task to study various cognitive processes including learning, memory, error monitoring, and response inhibition (20–28). This task has also been used to understand sensory processing and speed-accuracy trade-offs in rodents (3, 20, 21, 29). Modified versions like stop-signal tasks have also been used to study behavioral inhibition control (27, 28). These studies utilising the Go/No-Go task have used different sensory modalities such as olfaction (20–22), audition (30), and somatosensation (31–33). Behavioural readouts are extracted by monitoring the nose-poke, stimulus sampling or licking behavior (20–23, 25). Although seemingly simple, mouse licking behaviour shows nuanced motor patterns comparable to primate hand-reaching movements (34).

High-speed tongue imaging showed that successful licks contain sub-corrective movements while searching a spout, similar to primate hand-reaching. These corrections are context-dependent and governed by sensory feedback. Cue-evoked anticipatory licks have complex trajectories with long durations, while water-retrieval licks are more stereotyped (34). Further, rodent lick features help distinguish the mechanical act of consumption from cognitive and emotional states (35–38). The intra-burst inter-lick interval indicates task engagement and brainstem pattern generator integrity (35). Lick burst structure serves as a proxy for internal affective states, with mean burst size measuring palatability relative to reward concentration (38, 39). The number of bursts and inter-burst pause duration reflect motivation (37). This temporal architecture reveals emotional processes; truncated initial bursts can indicate anxiety (35, 36), while burst patterns can differentiate between innate and learned aversions (38). Taken together, monitoring lick behaviour in a Go/No-Go perceptual decision-making task can provide an insight into rodent perceptual decision making and change of mind.

We studied “change of mind” in mice using a Go/No-Go olfactory discrimination paradigm by monitoring the revisions in lick responses happening during a “No-go” trial. Erroneous lick responses for “No-go” cue, classified as the False Alarms, decreased gradually, but did not cease completely. These False Alarm trials exhibited variable licking patterns. We classified two False Alarm subpopulations: one showing revision of responses within trial by stopping of lick activity, resembling change of mind, as the revision trials (RT), and another with persistent licking similar to the correct hit trials, as non-revision trials (NRT). Based on the lick pattern analysis between RT and NRT, we conclude that mice exhibit response revision due to change of mind in a Go/No-Go olfactory discrimination task. Further, inhibition in OB is essential for improving the signal-to-noise ratio, hence sharpening the odor percept, and enhancing the relevant sensory signals over the noise (20, 22). Thus, manipulating GAD65-expressing intraneuronal activity could help understand the role of odor processing over RT and NRT. Therefore, we modulated the inhibitory network of OB during a complex odor-discrimination task to understand the role of olfactory bulb microcircuitry in shaping the revision behaviour.

## Results

### Mice revise the responses inappropriate to the context

To investigate whether mice revise context-inappropriate responses, we trained them in a Go/No-Go olfactory decision-making task under head-restrained conditions. Such a paradigm allows us to quantify commission and omission errors, along with correct responses, in the absence of any implemented punishments (Figure 1A). Mice learned a simple odor discrimination task quickly and accurately, reaching performance levels above 80%. (Figure 1B). The frequency distribution of the number of licks and total lick duration for the false alarm (FA) trials showed a skewness. Their distribution showed distinct subpopulations showing high and low lick numbers and durations (Figure 1C and D). Licking in a subset of false alarm trials was comparable to the lick number and total lick duration exhibited in correct hit trials, while the other subset had a low lick number and total lick duration (Figure 1C and D). Lick pattern varied within the False Alarm population, demarcating two types of commission errors, namely revision and non-revision trials. In revision trials, mice exhibited lick responses following stimulus onset and deliberate stopping, indicating error awareness leading to revision of the initial response. On the contrary, a persistent licking activity throughout the response window, similar to correct hit trials, was observed in the non-revision error trials (Figure 1E and F). The licking activity in both revision and non-revision trials started within 200-400ms of odor presentation. Animals revised their initial licking behaviour to stopping in revision trials, whereas non-revision trials exhibited continued licks till the odor offset (Figure 1G).

**Figure 1.**
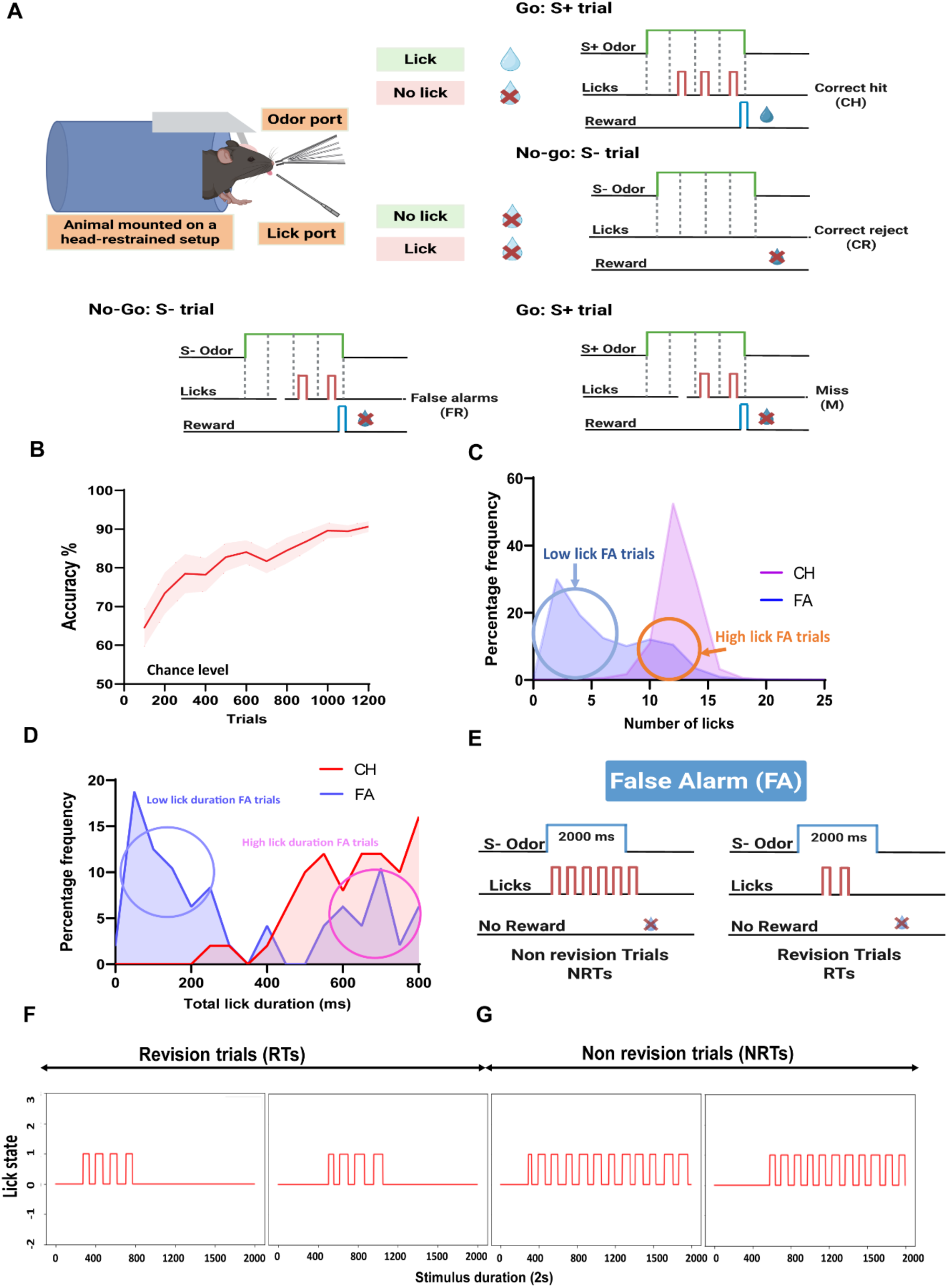
Mice revise their wrong decisions. A. Schematic illustration of the head-restrained olfactory Go/No-Go discrimination task, and types of trials registered based on the licking criteria. Odor duration is of 2 s, which overlaps with the response window. Odor duration is divided into 4 bins of 500 ms each. Lick criteria for rewarded Go-trials include a minimum of three licks in three out of four 500 ms bins, and a total licking duration of 240 ms. B. Line curve showing the learning progress (accuracy of performance, average ± SEM, SEM is represented by shaded area) in a simple odor discrimination task (Amyl acetate versus Ethyl butyrate, N=18 mice). C. Frequency distribution of the number of licks exhibited for the False Alarm (FA) and Correct Hit (CH) trials. The histogram illustrates a unimodal Gaussian-like peak for CH trials (purple), where animals consistently produced a high number of licks during rewarded stimuli. In contrast, FA trials (blue) exhibited a skewed distribution: one peak at low lick counts (low lick FA trials) and another at higher lick counts (high lick FA trials), suggesting the existence of two subpopulations within FA responses, those where the animal licked minimally and those where it licked vigorously despite no reward. D. Frequency distribution of the total lick duration exhibited for the False Alarm (FA) and Correct Hit (CH) trials, suggesting the existence of two subpopulations within FA responses, those where total lick duration is low, and the other with high total lick duration similar to CH. E. Illustration of FA trial subpopulations and trial structure. The schematic explains Non-Revision Trials (NRTs) and Revision Trials (RTs) among FA responses. In NRTs, animals encounter a non-rewarded S- (No-go) odor but still exhibit numerous licks very similar to those in rewarded trials, yet no reward is delivered. In RTs, animals encounter a non-rewarded S- (No-go) and produce fewer licks, wherein licks start a few ms after stimulus onset, but animals revise their licking behaviour by stopping, resulting in fewer licks. This division visually underscores the two classes of FA responses: high-lick NRTs and low-lick RTs. F. Representative trials: Revision Trials (RTs). The plots show lick events in the form of 0/1 binary output with 0 depicting no-lick and 1 lick activity as red traces for RTs, reflecting fewer lick events over the stimulus duration (2 s), consistent with the low-lick FA subpopulation. G. Representative trials: Non-Revision Trials (NRTs). The plots depict lick events as red traces in binary format, i.e., 0 depicts no lick event and 1 depicts a lick in NRTs, where multiple licks are observed throughout the stimulus window, characteristic of the high-lick FA subpopulation.

### Revision trials exhibit longer licks and inter-lick duration and are dependent on the learning phase

Revision trials differ from the licking activity in CH, NRT and Miss trials (MT) in lick bout duration and inter-lick interval, suggestive of uncertainty regarding the trial outcome. The average duration of each lick and inter-lick intervals were significantly higher in revision trials compared to CH and NRT (Figure 2A One-way ANOVA with Tukey’s multiple comparison test, p (CH-RT) < 0.0001, p (MT-RT) < 0.0001, p (RT-NRT) < 0.0001, p (CH-NRT) > 0.05, p (CH-MT) > 0.05 and 2B One-way ANOVA with Tukey’s multiple comparison test, p (RT-CH) < 0.001, p (RT-MT) < 0.001, p (RT-NRT) < 0.01, p (CH-NRT) > 0.05, p (CH-MT) > 0.05). This variation in the lick microstructure for RT implied a low-confidence hesitant licking followed by stopping. Stopping behaviour implies self-correction upon error-detection.

**Figure 2:**
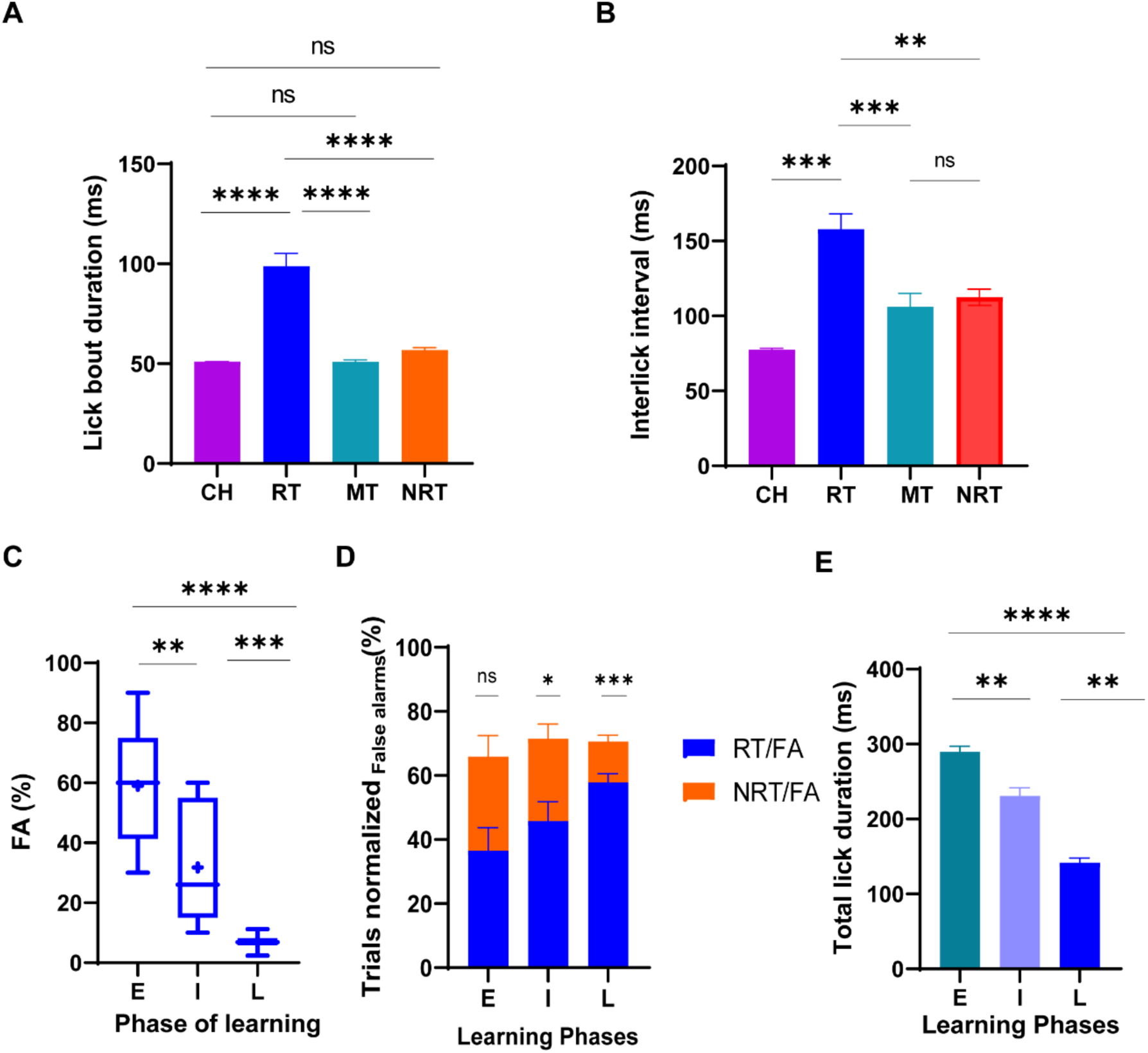
Learning phase-dependent revisions in a Go/No-go task. A Comparison of mean lick bout duration across trial types. The bar graph represents the average lick bout duration (ms) for CH, NRT, RT, and Miss trials (MT). Statistical analysis: One-way ANOVA with Tukey’s multiple comparison test, p (CH-RT) < 0.0001, p (M-RT) < 0.0001, p (RT-NRT) < 0.0001, p (CH-NRT) > 0.05, p (CH-MT) > 0.05, shows that RTs have significantly longer lick bout durations compared to other trial groups (****p < 0.0001), while differences between CH, NRT, and M are not significant (ns). B. Mean interlick interval for each trial type. The bar graph compares average interlick intervals (ms) across CH, RT, NRT, and MT trials. RT trials exhibit significantly longer interlick intervals compared to CH [One-way ANOVA with Tukey’s multiple comparison test, p (RT-CH) < 0.001, p (RT-MT) < 0.001, p (RT-NRT) < 0.01, p (CH-NRT) > 0.05, p (CH-MT) > 0.05, **p < 0.01, ***p < 0.001], N=18 for all the comparisons. C. Box-plot displays the FA percentage in the Early (E), Intermediate (I), and Learned (L) phases. One-way ANOVA followed by Tukey’s multiple comparison test detected highly significant differences: E vs I (p < 0.01, indicated by **), I vs L (p < 0.001, indicated by ***), and E vs L (p < 0.0001, indicated by **). D. Bar graph shows the percentage of Revision Trials (RT, blue) and Non-revision Trials (NRT, orange) normalized to the total number of False Alarm (FA) trials during Early (E), Intermediate (I), and Learned (L) training phases. The plot reveals an upward trend for RTs as learning progresses, indicating increased stopping behaviour and error monitoring in response to non-rewarded odors. Statistical comparisons were performed using one-way ANOVA followed by Tukey’s multiple comparisons test, p(I(RT/FA-NRT/FA)) <0.05, p(L(RT/FA-NRT/FA)) <0.001. Significant differences between phases are marked as follows: ns = not significant; * p < 0.05; *** p < 0.001. E. Mean total lick duration in RTs with increase in accuracy. Bar graph displays the mean ± SEM total lick duration (ms) during RTs for each learning phase (E, I, L). Data indicate that as the accuracy increases through training, total lick duration in RTs significantly decreases. One-way ANOVA with Tukey’s multiple comparisons test reveals highly significant differences, p(E-I)<0.01, p(I-L)<0.01, p(E-L)<0.0001, (** p < 0.01, **** p < 0.0001) among groups.

We hypothesised a learning-dependent increase in the revision trial frequency among the overall false alarm population, owing to better discrimination and error-detection. Learning stages were defined based on the performance accuracy as early learning (E, up to 60%), intermediate learning (I, >60<80%), and learned phases (L, ≥80%). As expected, false alarm frequency decreased as the learning progressed [Figure 2C, one-way ANOVA followed by Tukey’s multiple comparison test: p(E vs I) < 0.01, p(I vs L) < 0.001, and p(E vs L)< 0.0001]. This suggests a betterment in discrimination ability of the animals. We quantified the relative frequency of RT and NRT among FA population with learning. We could observe a linear increase in the RT frequency among FA population with the learning stages. Animals exhibited highest RT frequency at the learned stage [Figure 2D, one-way ANOVA followed by Tukey’s multiple comparisons test, p(I(RT/FA-NRT/FA)) <0.05, p(L(RT/FA-NRT/FA)) <0.001]. Moreover, we also quantified total lick duration within RT at different learning stages. We observed a consistent decrease in the total lick duration of RT with learning [Figure 2E, one-way ANOVA with Tukey’s multiple comparisons test, p(E-I)<0.01, p(I-L)<0.01, p(E-L)<0.0001]. This suggests that erroneous responses decrease with learning. These observations indicate that the error responses are likely to be followed by a revision in licking behaviour in the learned condition.

### Quick decision revisions happen in the learned stage

When we plotted the average lick probability of revision trials (RTs) population for individual animals performing with an accuracy more than 80%, a distinct temporal pattern emerged: animals tended to concentrate their licking within a specific early time window, most often reaching peak activity in the first 500–600 ms after the odor stimulus was presented. After this brief burst of licks, their responses rapidly diminished, indicating that they were actively stopping their behavior well before the end of the stimulus period. Examining the data on an animal-by-animal basis, we consistently observed this peak licking near stimulus onset, followed by stopping of the responses. (Figures 3A and B). This highlights both a shared strategy and individual nuances in how animals managed self-correction and revisions. To understand this at a broader scale, we pooled all RTs from multiple animals trained for the AA/EB odor discrimination task (Figure 3C) and compared their licking activity to the first 40 ms of stimulus presentation time (taken as baseline). The analysis revealed that, at a population level, the brief, early lick responses in RTs were significantly elevated over baseline activity, and ended rather quickly (Figure 3D). We were able to identify a time window by comparing licking activity (Figure 3D, paired two-tailed t-test with the baseline lick activity, adjusted alpha value with Bonferroni correction) during which the lick activity increased from the baseline, i.e., crossed the alpha value and returned back to the baseline. This time window was demarcated as the decision revision window (RW). (Figure 3D, RW, window duration demarcated as 286ms (demarcating lick initiation)-758ms (demarcating lick stopping)).

**Figure 3:**
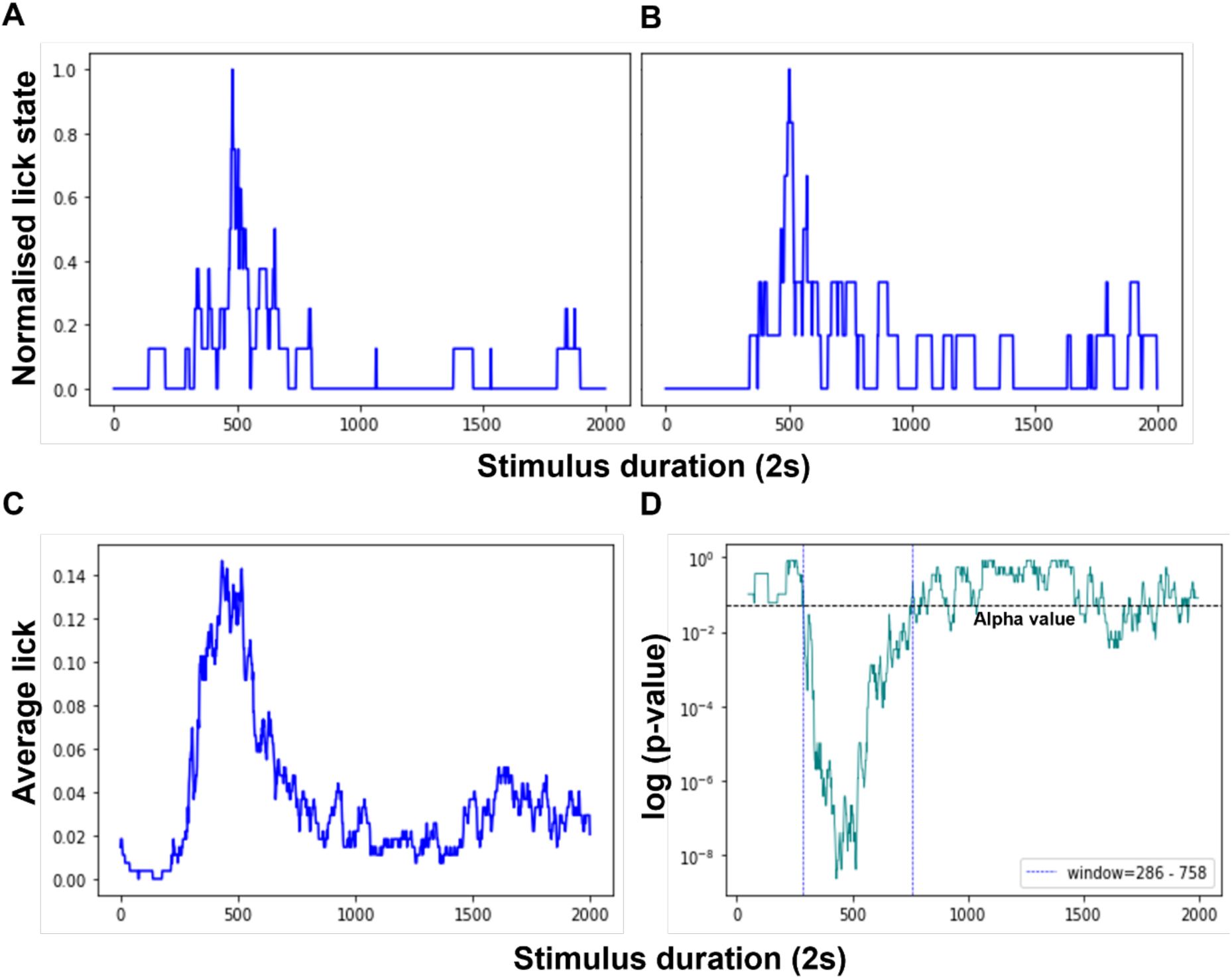
Quick decision revisions happen at the learned stage. A and B. Normalized lick probability for individual animals during Revision Trials (RTs) during the learned stage. Each panel presents the time-resolved lick probability curve for two different animals in the context of RTs (150-200 trials per animal). The x-axis represents stimulus duration, where 0 ms marks the onset of the odour stimulus and 2000 ms marks the end of the stimulus epoch. The y-axis reflects normalized lick state, demonstrating that both animals showed a prominent peak in licking activity within the first 500-550 ms after stimulus onset, followed by a rapid decline as the animals suppressed further licking during the remainder of the stimulus window, illustrating active stopping behavior. C. Population average lick probability for all RTs from a monomolecular odor discrimination task. This panel aggregates lick data across a group of animals performing an AA/EB monomolecular odor discrimination task (N=9), all RTs combined within the discrimination task group. A clear and consistent population-level peak occurs shortly after stimulus presentation, followed by a systematic reduction in licking, supporting the notion that RTs across animals and trials share a stereotyped temporal structure of brief responding followed by cessation. D. Statistical significance of licking changes during RTs (p-value time course). The p-value plot shows results from a two-tailed, paired t-test (compared with the baseline licking: first 40 ms after odor onset) with Bonferroni corrections. The horizontal dashed line marks the Bonferroni corrected alpha threshold for significance. P-values falling below alpha value early in the stimulus epoch indicated the initial surge in licking which is significantly elevated over baseline, while the return above alpha in the later part of the stimulus window identifies the point at which licking ceases and no longer differs from the baseline, quantifying both lick initiation and active stopping in RTs and the revision window marked by the time spanned during significant licking activity.

### Revisions facilitate latency adjustment and correct choice selection in subsequent trials

We further analysed how the occurrence of erroneous responses affects performance in subsequent trials. We quantified the probability of getting correct or incorrect rewarded (S+)/non-rewarded (S-) trials following RT, NRT, and MT. To understand this, we segregated animal-wise RTs, NRTs and MTs and recorded the latency, trial type (rewarded (S+) and non-rewarded (S-)), and trial outcome (PASS/FAIL) following them. We observed a higher probability of successful trials of both the rewarded and non-rewarded conditions, succeeding RT and NRT (Figure 4A, Paired t test, p>0.05, Figure 4C, Paired t test, p>0.05). However, P(correct)n+1S+ was at chance level right after the MT. Additionally, P(correct)n+1S- was very high for the trials succeeding MT (Figure 4B, Paired t test, p< 0.001). This suggests that the occurrence of RT and NRT led to higher probability of correct S+ as well as S-, while a decrease in the probability of correct S+ trial following MTs. This implies that MTs are a consequence of relatively lower motivation levels to engage in the task, unlike RTs and NRTs. We also quantified trial-wise lick latency in these (n+1)th S+correct trials succeeding RT, NRT and CH (the cases of 2 consecutive S+ correct trials) to judge if the animal adjusts its response latency to a faster or slower side following the erroneous judgements (either revised or not) in the preceding trial. Here, the case of latency adjustments for the second S+ correct during two consecutive S+ correct trials signalled how correct judgements and reward acquisition affected the latency in the subsequent trial. We observed that the animals significantly reduced their response latency for S+ trials following a correct S+ trial and tended to respond faster (Figure 4D, Paired t test, p< 0.05). However, for the S+ correct trials coming after the RT, animal’s mean lick latency increased significantly (Figure 4E, Paired t test, p< 0.05). Mean lick latency remained unchanged for the correct S+ trial following NRT (Figure 4F, Paired t test, p>0.05). This suggested that RTs are indeed trials wherein an animal monitors the error, revises the erroneous licking within stimulus presentation, and performs necessary response adjustment in the following trial to make better judgments. The delayed lick latency may signal higher vigilance and careful lick initiation instead of prompting. Lack of response adjustments following NRT could mean a continuous state of uncertainty, where animals are not able to judge their error and therefore no adjustments are made. This means that committing errors in anon-rewarded trial, specifically RT, enhanced animals’ ability to be more vigilant to monitor the error and respond more carefully in the subsequent trial.

**Figure 4:**
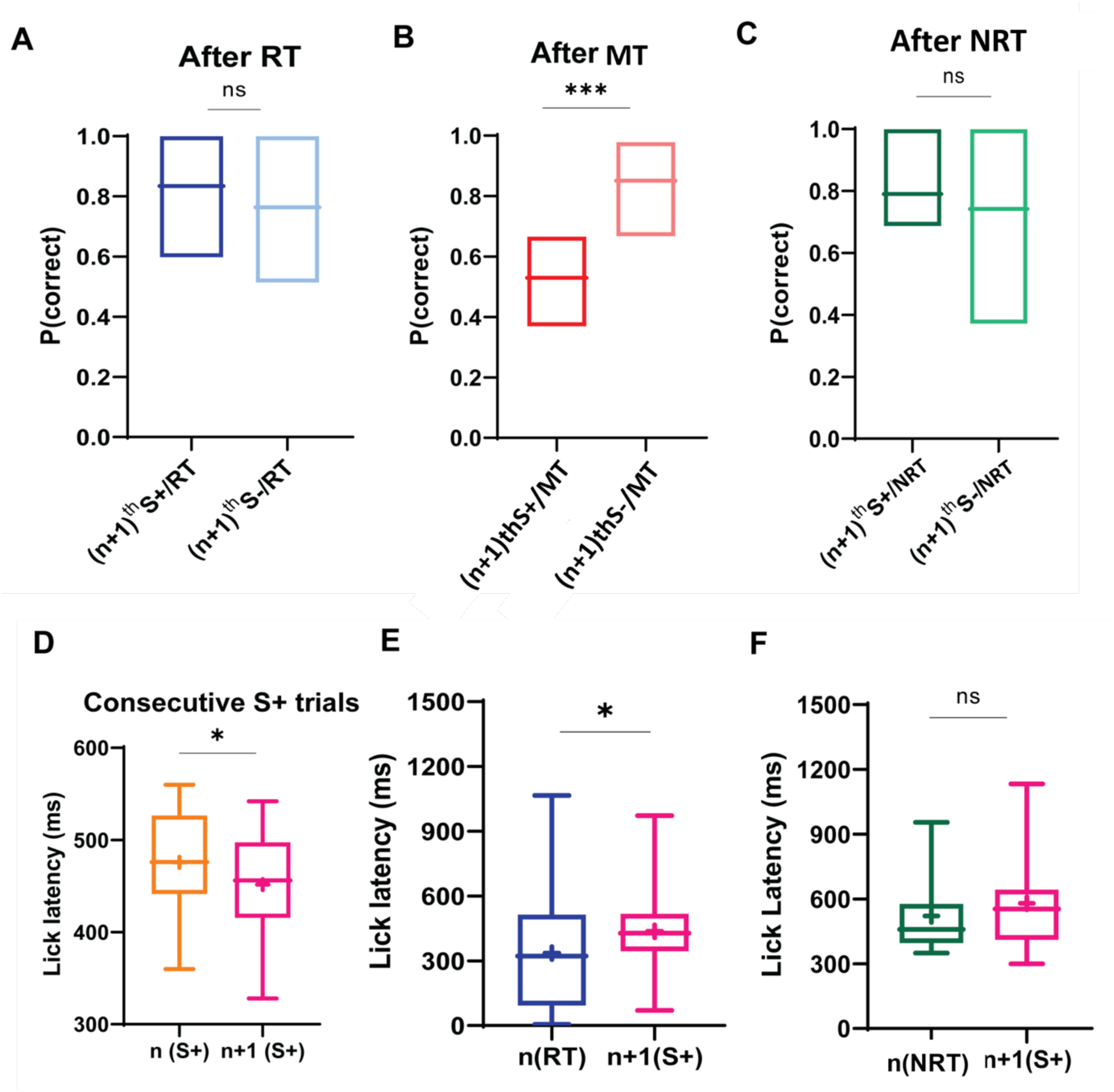
Occurrence of RT affects the subsequent trial outcome and response latency. A-C. Probability of performing a correct S+ or S- trial subsequent to RT, MT and NRT. Box plots show the probability of achieving a correct response for rewarded (S+) and non-rewarded (S-), depending on the immediately preceding trial type: A. After a revision trial (RT), the probability of a correct S+ and S- response did not significantly differ (Paired t test, p>0.05, ns). B. After a miss trial (MT), the probability of a correct S+ response was significantly lower compared to the S− response, indicating that misses could be arising due to low motivation of animals to engage in the task (Paired t test, p< 0.001***). C. After a non-revision trial (NRT), correct S+ and S- response probabilities did not show significant differences (Paired t test, p>0/05, ns). D-F. Lick response latency for consecutive rewarded trials. Box plots depict how the latency to lick on a correct rewarded trial (S+) changes as a function of the previous trial type D. Lick latency decreased significantly on the n+1^th^ (S+) trial compared to the preceding i.e. n^th^ S+ trial (Paired t test, p< 0.05*), suggesting that the correct hit trials lead to a decrease in response latency in the subsequent trial, if two rewarded trials were consequently presented. E. Following an RT, lick latency on the next S+ trial is significantly longer than in the preceding revision trial (Paired t test, p< 0.05*), indicating an adjustment in lick latency after revision trials, suggesting animals’ vigilance in responding more carefully. F. When the previous trial was NRT, lick latency on the next S+ trial does not significantly change (Paired t test, p > 0.05, ns), suggesting no major adjustments in response timing following non-revision trials. Statistical significance is indicated as follows: *p < 0.05; ***p < 0.001; ns, not significant

### Stimulus complexity alters the frequency and course of successful revisions

As reported earlier, binary mixtures evoke overlapping spatial patterns of neuronal activity in the dorsal olfactory bulb, leading to increased perceptual similarity and reduced discriminability (3, 20, 21). Binary mixture discrimination, a complex task, therefore carries a higher perceptual load. Hence, to elucidate if stimulus complexity affects revisions, we trained three groups of mice for monomolecular and binary odor mixtures with increasing complexity. For the complex mixture discrimination task, half of the animals received 60% AA + 40% EB or 55% AA + 45% EB as rewarded odor and 40% AA + 60% EB or 45% AA + 55% EB as non-rewarded odors and vice-versa. All odor mixtures were 1% v/v diluted in 5mL of mineral oil. During early learning phases the occurrence of false alarm trials was similar across the complexity group [Figure 5B, One-way ANOA, Tukey’s multiple comparisons test p(AA/EB-AA/EB 60/40)>0.05, p(AA/EB-AA/EB 55/45)>0.05, p(AA/EB 60/40-AA/EB 55/45)>0.05]. Revision trials (RTs) were highest in the simple odor discrimination group compared to the highly complex [Figure 5C, One-way ANOVA, Tukey’s multiple comparisons p(AA/EB-AA/EB 60/40)>0.05, p(AA/EB-AA/EB 55/45) <0.01]. Non-revision trials (NRTs) were highest for complex odor mixture discrimination compared to the simple odor discrimination [Figure 5D, One-way ANOVA, Tukey’s multiple comparisons test, p(AA/EB-AA/EB 60/40)<0.01, p(AA/EB 60/40-AA/EB 55/45)>0.05, p(AA/EB-AA/EB 55/45)<0.01]. This suggests poor discrimination and error detection for complex discrimination tasks at the early learning stages. As expected, increase in complexity resulted in higher false alarms in the learned phases [Figure 5E, One-way ANOVA, Tukey’s multiple comparisons test, p(AA/EB-AA/EB 60/40)<0.0001, p(AA/EB 60/40-AA/EB 55/45)<0.05, p(AA/EB-AA/EB 55/45)<0.0001].

**Figure 5:**
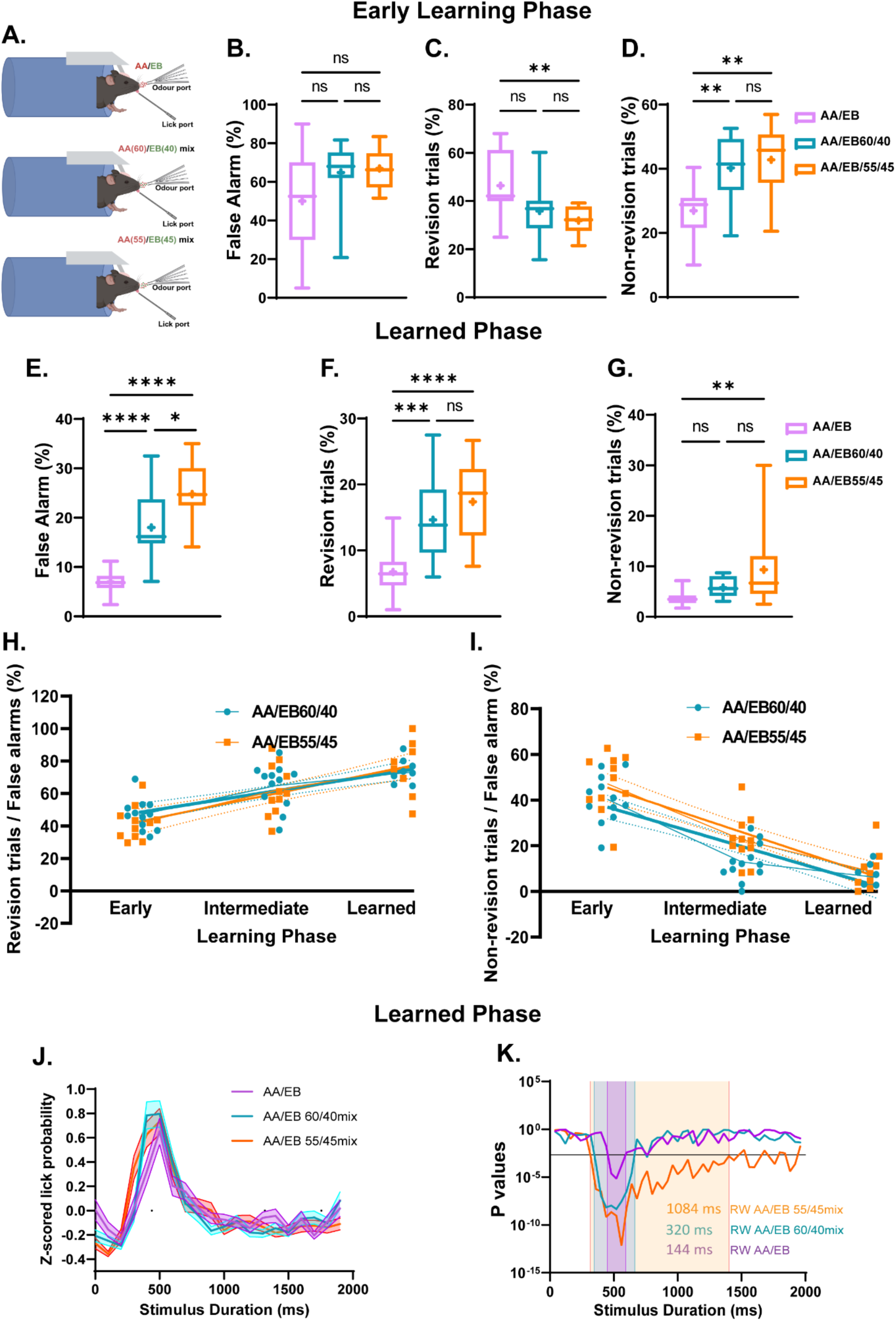
Stimulus complexity alters the frequency and duration of revisions in a learning phase-dependent manner. A. illustration depicting Go/No-go behavioral paradigm under head-restrained conditions. B. B-D. Early learning phase: B. False Alarm frequency did not differ significantly across complexity levels, indicating similar early error rates regardless of perceptual complexity. [One-way ANOA, Tukey’s multiple comparisons test p(AA/EB-AAEB60/40)>0.05, p(AA/EB-AA/EB/55/45)>0.05, p(AA/EB60/40-AA/EB/55/45)>0.05] C. Revision trial (RT) percentage is significantly lower for the AA/EB monomolecular odor discrimination compared to the complex mixture [One-way ANOVA, Tukey’s multiple comparisons p(AA/EB-AA/EB60/40)>0.05, p(AA/EB-AA/EB55/45) <0.01]. D. Non-revision trial (NRT) percentage is significantly higher with increasing complexity, suggesting diminished error awareness with more complex stimuli [One-way ANOVA, Tukey’s multiple comparisons test, p(AA/EB-AA/EB60/40)<0.01, p(AA/EB60/40-AA/EB/55/45)>0.05, p(AA/EB-AA/EB/55/45)<0.01]. E-G Learned phase: E. Box plots showing a comparison of False Alarm frequency across different complexity in learned state. FA% are higher in complex binary discrimination task compared to monomolecular discrimination, reflecting persistent perceptual ambiguity in difficult conditions [One-way ANOVA, Tukey’s multiple comparisons test, p(AA/EB-AA/EB60/40)<0.0001, p(AA/EB60/40-AA/EB/55/45)<0.05, p(AA/EB-AA/EB/55/45)<0.0001]. F. Revision (RT) frequencies mirrored the trend of high occurrence in the complex mixture discrimination compared to the simple odor discriminations largely due to the decrease in the overall False Alarm population [One-way ANOVA, Tukey’s multiple comparisons test, p(AA/EB-AA/EB60/40)<0.0001, p(AA/EB60/40-AA/EB/55/45)<0.05, p(AA/EB-AA/EB/55/45)<0.0001], indicating the influence of stimulus complexity. G. Non-revision trials (NRT) showed a similar trend of increasing frequency with the complexity [One-way ANOVA, Tukey’s multiple comparisons test, p(AA/EB-AA/EB60/40)>0.05, p(AA/EB60/40-AA/EB/55/45)>0.05, p(AA/EB-AA/EB/55/45)<0.01]. All data represent box plots (min-to-max whiskers, horizontal line for median, ‘+’ within the box represents mean). ns. denotes non-significant, *p < 0.05, **p < 0.01, ***p < 0.001, ****p < 0.0001 by one-way ANOVA with Tukey’s multiple comparisons test. H. Fraction of revision trials (RT), normalized to total false alarms, plotted across early, intermediate, and learned phases for both AA/EB60/40 and AA/EB55/45 odor mixtures. RT frequency shows a linear increase with learning for both complexities (r² = 0.6 for AA/EB60/40; r² = 0.5 for AA/EB55/45; p < 0.05). I. Fraction of non-revision trials (NRT), normalized to total false alarms, assessed across the same learning stages and mixture complexities. NRT frequency decreases with learning, reflecting reduced uncorrected errors as mice become proficient with both AA/EB60/40 and AA/EB55/45 odor mixtures. J. Z-scored lick probability traces exhibited by mice in the learned phase for three odor discrimination tasks with increasing complexity (AA/EB, AA/EB60/40 mixture, AA/EB55/45 mixture). K. P-value plot for licking behavior during revision trials (RT) at the learned stage across different odor complexities. Shaded regions indicate the defined Revision windows for odor pair discrimination task. Statistical comparison used paired t-test; alpha value set at 0.001 (Bonferroni-corrected).

Consequently, revision and non-revision trials were also higher for the complex odor discrimination compared to the simple AA/EB discrimination group [Figure 5F, One-way ANOVA, Tukey’s multiple comparisons test, p(AA/EB-AA/EB 60/40)<0.0001, p(AA/EB 60/40-AA/EB 55/45)<0.05, p(AA/EB-AA/EB 55/45)<0.0001 and 5G, One-way ANOVA, Tukey’s multiple comparisons test, p(AA/EB-AA/EB 60/40)>0.05, p(AA/EB 60/40-AA/EB 55/45)>0.05, p(AA/EB-AA/EB 55/45)<0.01], as these two are the two subsets of false alarm population. However, an interesting pattern emerged when we quantified the relative dominance of RTs and NRTs in the total false alarm population. We observed a linear increase in the false alarm normalised RT percentage with learning in both AA/EB 60/40 and AA/EB 55/45 complex mixture discrimination [Figure 5H, Rsquared = 0.68 (AA/EB 55/65), 0.54 (AA/EB 60/40), p <0.001]. Subsequently, a linear decrease in false alarm normalised NRTs was observed with learning (Figure 5I). This suggests that as learning happens, revisions become more frequent irrespective of stimulus complexity.

Additionally, we observed fewer licks in the entire duration of the response window with the simple monomolecular odor discriminations compared to binary mixtures for the commission error trials. RT lick pattern yielded a sharp initiation with a single peak at around 500-600ms irrespective of the complexity. However, the reversals took longer for complex odors, compared to monomolecular ones (Figure 5I & K). In summary, the reversal window duration was decided by the complexity of the task. RW was progressively longer for the complex and highly complex odor discrimination tasks compared to monomolecular discriminations (Figure 5K). These results suggest that the stimulus complexity alters revision behavior, especially the stopping behavior, which could arise due to uncertainty.

### Optogenetic activation of olfactory bulb GAD65 inhibitory interneurons modulates revisions

Optogenetic activation of OB GAD65-expressing intraneuronal population facilitated learning and discrimination ability of animals in a complex odor discrimination task (Figure 6 B-D). Consistently higher D-prime values of GAD65-ChR2 group compared to the GAD65-EYFP controls suggest that GAD65-ChR2 groups were better able to distinguish between the S+ and S- odors (Figure 6B). GAD65-ChR2 exhibited higher frequency of correct hits (CH%) during early (E) and intermediate (I) learning stages [Figure 6C, Two-way repeated measure ANOVA, Sidak’s multiple comparisons test p(E(GAD65-EYFP-GAD65-ChR2)) and (I(GAD65-EYFP-GAD65-ChR2))<0.05, ns = not significant]. GAD65-ChR2 groups were faster in discriminating the odors during the learned phase compared to GAD65-EYFP (Figure 6D, Unpaired Two-tailed t test, p(GAD65-EYFP-GAD65-ChR2) < 0.01), suggesting better discrimination ability upon increased inhibition in the OB. In order to examine whether increased inhibition affects revision phenomena, we quantified the percentage occurrence of revision trials (RT%) in GAD65-ChR2 and GAD65-EYFP (control) groups in the learned stage i.e. when both the groups performed with criterion level accuracy (80% and higher) [Figure 6E, Unpaired Two-tailed t test p(GAD65-EYFP-GAD65-ChR2) < 0.05]. We found a reduction in overall error responses in both groups upon learning. RT percentage was significantly lower in GAD65-ChR2 group compared to GAD-EYFP group, suggestive of enhanced perceptual disambiguity and increased error awareness upon increasing inhibition in OB (Figure 6E). We also quantified percent decrease in the RT and NRT trials animal-wise from the early learning phase to the learned phases [Figure 6F, Two-way ANOVA with Tukey’s multiple comparison test; *p(RT(GAD65-EYFP-GAD65-ChR2) <0.05 and p(NRT(GAD65-EYFP-GAD65-ChR2) <0.05. Error bars represent SEM]. Percentage decrease was quantified by subtracting the number of animal-wise RT and NRT exhibited in learned condition from the early learning phase (E) and normalising it with the same. This indicated a higher percentage reduction in RT and NRT trials with learning in GAD65-ChR2 animals compared to GAD65-EYFP group, indicating that the overall error rate decreased with the increased inhibition in olfactory bulb. Additionally, GAD65-EYFP animals exhibited an extended revision-window compared to GAD65-ChR2 animals, indicating longer lick duration in controls compared to groups with increased inhibition in OB (Figure 6H). GAD65-ChR2 group showed higher percentage reduction in RT and NRT in the learned stages compared to the control group, suggesting a betterment in error monitoring and rectification in groups with increased inhibition (Figure 6F).

**Figure 6:**
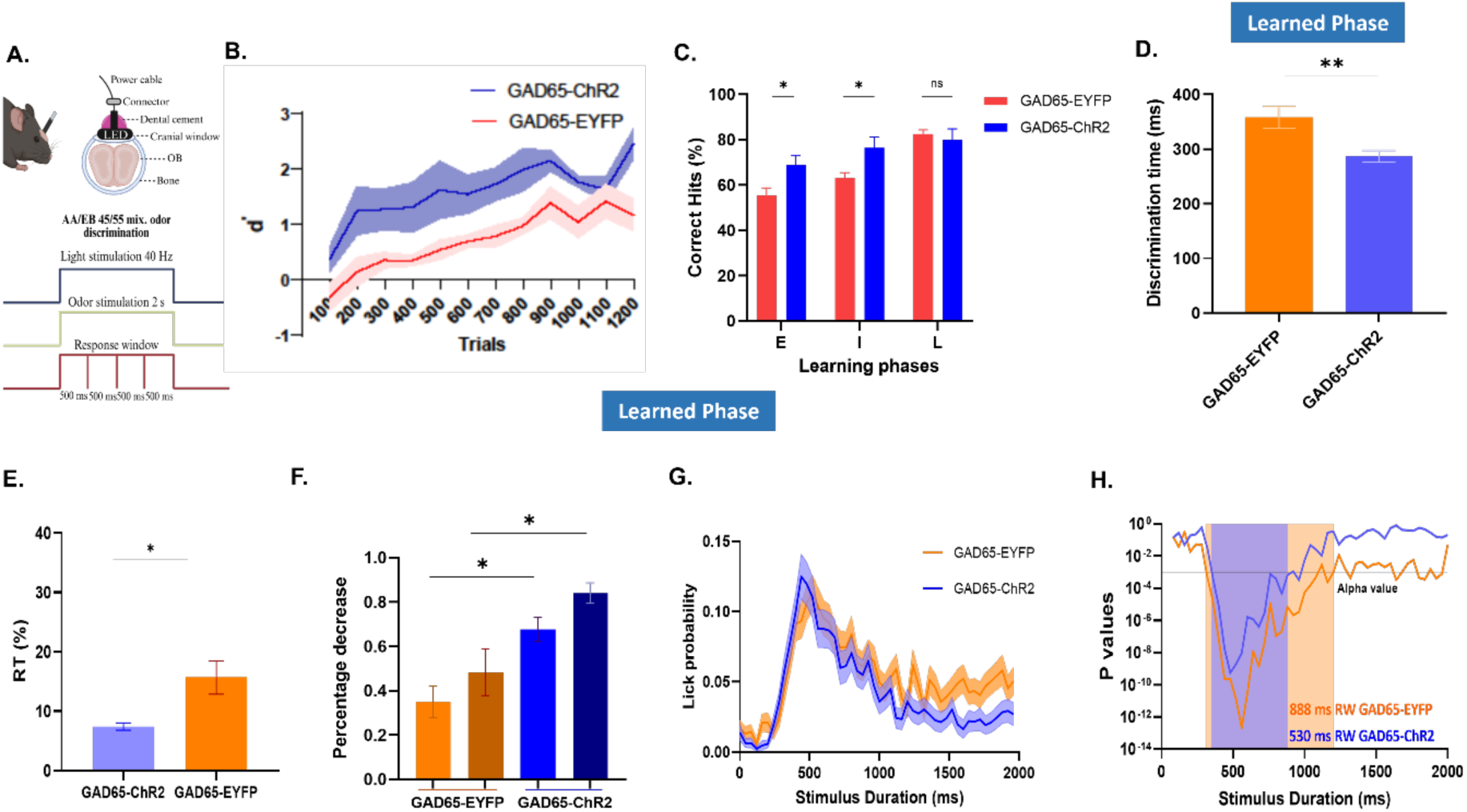
Optogenetic activation of olfactory bulb GAD65 inhibitory interneurons modulates revisions. A. Schematic illustrating cranial window preparation and LED implantation for optogenetic stimulation in mice in the upper panel, and experimental timeline showing odor stimulation (2 s), concurrent light stimulation (40 Hz), and response window segmentation for behavioural trials in the lower panel. B. d′ (d-prime) values across trials for GAD65-EYFP and GAD65-ChR2 groups, indicating odor discrimination sensitivity during learning. Shaded regions indicate SEM. C. Bar-graph showing significantly higher correct hit (CH%) exhibited by GAD65-ChR2 than GAD65-EYFP groups at different accuracy levels. GAD65-EYFP mice catch up at the higher accuracies i.e. in the learned stage (accuracy 80%). Statistical comparison done using Two-way repeated measure ANOVA followed by Sidak’s multiple comparison test, *p [E(GAD65-EYFP-GAD65-ChR2)] and [I(GAD65-EYFP-GAD65-ChR2)]<0.05, ns = not significant, N_GAD65-EYFP_=14, N_GAD65-ChR2_ = 13. D. Average discrimination time (ms) taken by animals in GAD65-EYFP and GAD65-ChR2 groups to distinguish between odors of high complexity (AA/EB 55/45 mix) during the learned phase. GAD65-ChR2 mice showed significantly faster discrimination times. Statistical analysis was performed using Unpaired Two-tailed t-test [**p(GAD65-EYFP-GAD65-ChR2) < 0.01]. E. Percentage of revision trials (RT) for GAD65-EYFP and GAD-ChR2 groups during the learned phase, analysed using unpaired t-test; *p(GAD65-EYFP-GAD65-ChR2) < 0.05. F. Percent decrease in RT and NRT from early learning (E) to learned phase (L) in GAD65-EYFP and GAD65-ChR2 groups, analysed using Two-way ANOVA with Tukey’s multiple comparison test; *p(RT(GAD65-EYFP-GAD65-ChR2) <0.05 and p(NRT(GAD65-EYFP-GAD65-ChR2) <0.05. Error bars represent SEM. G. Lick probability traces for all the RT combined in the learned phase for GAD65-EYFP and GAD65-ChR2 groups; shaded areas represent SEM. H. Logarithmic p-value plot across stimulus durations for both groups, with shaded regions highlighting revision windows for GAD65-EYFP and GAD65-ChR2; paired two-tailed T-test, statistical threshold indicated by Bonferroni adjusted alpha value (p=0.001), N_GAD65-EYFP_=14, N_GAD65-ChR2_ = 13

## Discussion

### Mice exhibit response-revisions in an olfactory Go/No-Go discrimination task

As mice learned the olfactory Go/No-Go discrimination task, they displayed nuanced behavioural flexibility, especially in how they responded to non-rewarded odors. Current literature characterises this responses to non-rewarded odors as wrong ones, ignoring the nuances of this behaviour (26, 40–42). Lick pattern analysis revealed two distinct subpopulations among false alarms: non-revision trials (NRT), where the lick behaviour was incessant and near-indistinguishable from rewarded correct hits (CH), and revision trials (RT), marked by fewer licks and earlier lick cessation. In revision trials, mice exhibited a behavioural fingerprint of proactive error correction (27). This ability to deliberately revise an ongoing action is thought to reflect underlying confidence in judgments and real-time error monitoring. Previous reports found that mice employ a combination of two decision-making strategies: at times, they act swiftly due to a sense of urgency, while at other times, they continue to integrate new sensory information even after action initiation. The “Parallel Sensory Integration and Action Model,” captures the idea of response updating with continuous sensory integration (43). This indicates that mice possess the ability to adjust their actions dynamically, responding to new information in real-time rather than relying solely on their initial behavior (26, 43).

The temporal structure of the lick pattern in the revision trials (RT) is brief, stimulus-driven initiation followed by cessation, suggesting that animals are not just committing the mistakes but are actively detecting and correcting them, underscoring the presence of higher cognitive and executive control and trial-by-trial confidence estimation. This finding adds more weight to the idea that mice can be aware of their own actions and revise responses when facing uncertainty or conflicting evidence. Perceptual decision-making is not always a single, instantaneous act. Sensory integration can be carried out in parallel with the behavioural responses (4, 6, 7, 44), implying that animals may start responding and then alter their action if later evidence implies an error or uncertainty (45). The concept of revision windows defined in the present study is the period where behaviour is flexibly updated post-initiation. Based on the existing literature around the flexibility allowing response update and sensory integration post-action initiation (5–7, 45–47), we argue that the revision window (RW) represents an integral part of perceptual decision-making wherein responses are flexibly modulated based on evidence accumulation, leading to a better odor percept. Thus, our results illustrate a general strategy adapted by mice in a basic olfactory Go/No-Go discrimination task.

### Do decision revisions facilitate correct choice selection in subsequent trials?

Recent findings highlight a fascinating aspect of animal behaviour, i.e., making and recognizing errors in a task, can act like a task reset and boost accuracy (46–49). In our data, following a revision trial (when mice started to lick and then rectified their licking mid-trial by stopping), or a non-revision trial (when it continued with licking response despite a non-rewarded odor), the probability of subsequent trials to be correct irrespective of being a “Go” or “No-go” trial suggests that the occurrence of RT and NRT affect the performance accuracy positively. The latency adjustment following RT suggests that committing an erroneous action makes animals more cautious and attentive for the subsequent trials (Figure 4-E). A similar phenomenon has been studied in the context of an auditory discrimination task performed by rats, where rats showed post-error slowing of reaction, which is shown to be controlled by the dorso-medial Pre-frontal cortex (dmPFC) (47). Studies in rodents as well as in humans have shown that committing and monitoring errors help subjects re-engage with the task, recalibrate their attention, and fine-tune timing, improving performance in subsequent trials (4, 6, 7, 45–47).

This effect of improved tendency to perform the trial correctly is non-existent in trials succeeding a missed trial (MT, rewarded trials where animals failed to fulfil the lick criteria), where animals fail to engage in the task (Figure 4 B). Therefore, we argue that miss trials (MT) happen due to the motivation failure. Together, these results demonstrate that mice are able to self-monitor their errors, revise, and adjust latencies to the subsequent trials instead of responding hastily (Figures 3 and 4).

Furthermore, we could observe an experience-dependent change in the error monitoring and revision behaviour as mice progressed with the task (Figure 2C and D). Early on, mice showed more erroneous responses, but as their ability to discriminate between S+ and S-odors improved with training, the overall rate of false alarms (FA) declined, showing that learning enhanced their capacity to distinguish rewarded from non-rewarded odors (Figure 2C). Notably, the false alarm (FA) population underwent major reshaping with learning. At first, the FA population exhibited an equal share of RT and NRT. With learning, a bigger share of FA trials switched to revision trials (RTs), where the mice started to lick but then quickly stopped, actively correcting themselves with each trial. The decline in NRT with learning highlights the importance of enhanced perceptual clarity with experience (Figure 2D). Therefore, flexible error-monitoring and revisions emerged with learning. This notion could be supported by the line of studies investigating how perceptual decision-making and error-rates evolve with learning (48–51)

### Odor complexity imposes high perceptual load, elevating revisions

Odor complexity imposes a high perceptual load on mice, revealing clear effects on both the learning pace and the ability to discriminate between the rewarded and non-rewarded odor (3, 20, 21). Our results show that discriminating between complex mixtures such as AA/EB60/40 and AA/EB55/45 required substantially higher time compared to distinguishing simple monomolecular odors (AA/EB) (Figure 5C). This is consistent with a previous study from the lab demonstrating that similar odor mixtures evoke overlapping activity patterns in the olfactory bulb, making fine discrimination more challenging (3, 21). As a result, mice learning to discriminate between these mixtures showed a slower learning pace and required longer to reach the criterion accuracy (Figure 5A and B).

This increased perceptual load shapes the frequency and nature of revision and non-revision trials during the task (Figure 5B-G). In the early stages of training, all groups, regardless of odor complexity, exhibited high false alarm rates (Figure 5B), indicating poor initial discriminability and error awareness. However, as mice learned the discrimination task involving varying complexity, a striking difference emerged. The extent of learning was kept similar across the odor set to have a better comparison between simple and complex odor conditions (binary mixtures). We compared the revision and non-revision trials for the high-performance accuracy blocks across stimulus complexity groups. Even in the learned state, animals challenged with complex mixtures have persistently higher false alarm frequencies populated by a higher percentage of non-revision trials (NRTs), exhibiting sustained incorrect licking behaviour (Figure 5E and G). Meanwhile, the relative proportion of revision trials (RTs) among FA, rises linearly with experience, even for the complex discrimination tasks (Figure 5H). This suggests that animals commit more errors but become better in error monitoring and behavioral revision, particularly when perceptual ambiguity is high.

We further observed that the time window in which revisions occur, i.e, the revision window (RW), was delayed for complex mixture discrimination compared to simple odor discrimination (Figure 5J). This means that under higher perceptual load, the animal’s internal process for detecting and correcting mistaken actions takes longer to resolve, likely reflecting the difficulty in extracting clear sensory evidence from overlapping input. These observations highlight findings from perceptual integration models demonstrating that ongoing sensory processing and evidence accumulation can continue to alter actions after response initiation, especially when stimuli are ambiguous (43). Therefore, we conclude that stimulus complexity not only slows down learning but also elevates the demand for flexible, adaptive error monitoring and revision, highlighting the interplay between sensory processing and executive functions in decision-making.

### Increase in synaptic inhibition in the OB circuits affects stimulus-processing and thus revisions

Increasing inhibitory control within olfactory bulb (OB) circuits, through optogenetic activation of GAD65-expressing interneurons, has a pronounced impact on how mice process odors and adaptively monitor errors. Our results show that increasing OB inhibition via ChR2 activation in GAD65 neurons improved both the speed and accuracy of odor discrimination when mice are challenged with a highly overlapping mixture of odor stimuli (Figure 6B-D). Mice in the GAD65-ChR2 group consistently reached higher d-prime (d’) values and exhibited faster discrimination times than EYFP controls, indicating more precise sensory processing (Figure 6B and D). This enhancement was evident even in early and intermediate learning phases, where GAD65-ChR2 animals produced more correct hits, while controls took longer to catch up as accuracy increased (Figure 6C).

Enhanced inhibitory signalling is known to refine fine odor discrimination (20, 22, 23, 25). In the present study, we show that enhanced inhibition in the OB circuit affects error monitoring and revision behaviour as well. In the learned stage, both GAD65-ChR2 and GAD65-EYFP groups made fewer mistakes, but the GAD65-ChR2 mice showed significantly fewer revision trials (RTs) compared to controls (Figure 6E). This suggests that increased inhibition leads to better perceptual processing between rewarded and non-rewarded odors, reducing ambiguity and errors in general. In addition, the revision window, i.e., the temporal window in which animals could revise their erroneous responses, was shorter in the GAD65-ChR2 group, indicating that mice with stronger OB inhibition revised their responses less often but more rapidly (Figure 6H). The percent decrease in RT and NRT from the early learning phase to learned stages was also greater in the GAD65-ChR2 group, further supporting better error monitoring, due to lower perceptual ambiguity, and more time-efficient correction with enhanced OB inhibition (Figure 6F).

Collectively, these findings are in agreement with the studies showing that inhibitory interneurons in the OB shape sensory representations of the highly overlapping cues and limit aberrant behavioral responses. Therefore, this study highlights the essential role of OB circuits in coordinating higher-order cognitive processes with fine perceptual discrimination. Moreover, the simple design of the Go/No-Go paradigm has enabled its widespread use to understand different aspects of sensory-cognitive processes such as perceptual decision-making, working memory, and behavioral adaptability across humans, non-human primates, and rodents (3, 20, 21, 52–55). This has been widely used to evaluate sensory-cognitive and behavioural impairments in different neuropsychiatric and substance use disorders, especially in patients with ADHD and schizophrenia who exhibit increased response to No-Go stimuli and difficulty detecting stimuli (56–63). In addiction patients, the task assesses relapse risk, with electrophysiological measures predicting treatment outcomes (64). Therefore, a thorough study of the response variability observed and its attributes in a Go/No-Go task is required. In the present study, we thoroughly characterised and established a behavioural phenotype exhibited by animals that, to the best of our knowledge, had not been reported earlier. This behaviour is crucial in understanding the nuances of the responses and investigating the neural correlates underlying the change of mind.

## Materials and Methods

### Animals used

A total of 67 adult C57BL/6J male mice, aged 9-12 weeks old, were used for the Go/No-Go olfactory discrimination experiments. For optogenetic modulation of Olfactory bulb (OB) inhibitory neurons, i.e., GAD65 interneuron, the following animal lines obtained from Jackson’s laboratory were utilised:

B6N.Cg-Gad2^tm2(cre)Zjh/J^ (GAD65-cre)

B6.Cg-Gt(ROSA)26Sor^tm32(CAGCOP4*H134R/EYFP)Hze/J^ (ChR2-floxed)

B6.129X1-Gt(ROSA)26Sor^tm1(EYFP)Cos/J^ (EYFP-floxed)

GAD65-cre animals were crossed with ChR2-floxed and EYFP-floxed animals to generate GAD65-ChR2, and GAD65-EYFP animals, to express ChR2 and EYFP specifically in GAD65-expressing cells for the optogenetic experiments. A total of 13 GAD65-ChR2 and 14 GAD65-EYFP mice were used for the optogenetic study.

### Maintenance of animals used in the study

Animals were housed in individually ventilated cages (IVC) of floor area of 800 cm^2^, equipped with the air control unit in a 12 hrs dark-light cycle. Each cage housed 3-5 mice. They were provided with standard rodent bedding and nesting material. The holding room temperature was maintained at 22-25 °C, and the relative humidity at 45-55 %. All behavioural experiments were performed during the light cycle. Mice had ad libitum access to food but were on a restricted water schedule, ensuring that the body weight never fell beyond 80% of their initial body weight. The water restriction schedule never extended beyond 12 hrs.

### Ethical approval

The animal usage protocol used in this part of the study is approved by the Institutional Animal Ethics Committee (IAEC) at IISER Pune, and the Committee for the Purpose of Control and Supervision of Experiments on Animals (CPCSEA), Government of India (animal facility CPCSEA registration number 1496/GO/ReBi/S/11/CPCSEA). The animal usage protocol number are licensed under IISER/IEAC/2020_1_14.

### Surgical procedures Head-post implantation

Mice were anaesthetised with a cocktail of ketamine-xylazine (50 and 10 μg/g, IP). Depth of anaesthesia was confirmed by the absence of the toe-pinch reflex before mounting the animal on the stereotaxic apparatus. Eyes were moistened with artificial tears throughout the surgery. Lignocaine gel was applied locally to the skin region where an incision was required. The scalp was removed after mounting. The exposed skull was cleaned with ACSF (125 mM NaCl, 5 mM KCl, 10 mM glucose, 10 mM HEPES, 2 mM CaCl₂, 2 mM MgSO₄, pH 7.4), dried, and applied with Ivoclar Vivadent EcoEtch® for 20 s. Further, the skull was cleaned, dried, and applied with a UV-curing primer. UV-curing dental cement was applied on the primer after drying. UV-curing dental cement was also applied to the base of the metal head-post and affixed on the skull. Remaining exposed areas were covered with acrylic dental cement to ensure long-term stability. Mild bleeding was controlled using ACSF-soaked Gelfoam. Mice were kept on a heating pad until recovery, given moistened food, and monitored for 2 days, with dexamethasone (5 μg/g, in water) administered if inflammation occurred.

### Cranial window and LED implantation for optogenetic experiments

The headpost-implanted anaesthetised mouse was mounted on the stereotactic instrument. The skin overlaying the OB was removed using a blade and scissors. The connective tissue layer was scraped using forceps. The musculature surrounding the eye was cut carefully, while preserving the blood vessels. After cleaning with ACSF, a 2.5 mm biopsy punch was placed on the skull above the OB center, avoiding the infra-cerebral vein positioned caudally at the junction of the olfactory bulb and cortex. It was gently pressed and rotated to carve out a groove on the skull, which was augmented using a hand-held drill. Once thinned, the central skull piece within the groove was tested for flexibility and lifted using forceps and needle without damaging the dura. ACSF-soaked Gelfoam pieces (AbGel, SGK labs, Mumbai) were used to control bleeding. Dexamethasone was applied to prevent inflammation. A glass coverslip was placed on the exposed circular area with the dura beneath. Dental acrylic cement sealed the window and the remaining exposed skull area. If the window remained clear for 4-6 days post-surgery, a 2 mm LED (either blue: NFSB036BT or orange: NJSA172, Nichia Corporation) with a header-socket arrangement (ED11100-ND, ED8250-ND, Digikey) was fixed above the glass coverslip using dental cement. The animal was then removed from the stereotaxic apparatus and kept on the heating pad, and fed with moistened food pellets until recovery.

### Olfactory Go/No-Go discrimination task Paradigm

Each mouse was mounted in a PVC tube with a plexiglass platform wrapped in metallic mesh for the electrical connectivity during licking. Olfactory discrimination tasks were performed using a custom-built 10-channel olfactometer integrated with a lickometer, as described in previous studies (21, 23, 25, 29, 53, 65). The lickometer recorded and digitized lick responses toward odorants. The olfactometer was equipped with mass flow controllers and solenoid valves for precise odor delivery. A manifold splits clean air to different MFCs, with air through mini MFCs entering odor bottles, while the main MFC air served as a dilution stream. The streams merged at the output through a T-tube before entering the seven-channel glass delivery nozzle with an exhaust outlet. During the preloading time of 3.2 s, odorized air passed through the nozzle and was diverted to the exhaust until a steady state. The exhaust was switched off during the entire odor presentation time. Lick state 1 or 0 signalling lick or no-lick were timestamped and recorded with millisecond speed in a CSV file.

### Task habituation

Animals were not restrained on the first day and allowed to roam freely on the setup. Pretraining phases began the next day with a 20 trials session, each trial starting with a 200 ms tone immediately followed by a 3 μl water drop. The delay in water delivery following the tone was increased to 1 s for the next 30 trials in subsequent sessions and further to 2 s for the next 30 trials, training mice to wait patiently after the tone cue. Once the mice had mastered the waiting and licked reliably to retrieve water, a 2 s pulse of the odor (1% v/v in mineral oil) was introduced, and animals were rewarded for licking during the odor presentation. The licking criteria were made stringent successively as mice learned to lick during the stimulus presentation, from 80 ms one lick in the starting stages to a total 240 ms lick duration and 3 licks in three subsequent bins of 500 ms each during 2 s odor presentation time in the final stage. Water delivery was decided based on the licking criteria (240 ms total lick duration & at least one lick eachin three out of four 500 ms time bins during a 2 s of odor presentation time) during odor presentation, provided that the baseline licking (licking during 1 s before stimulus presentation) was avoided. Licking during baseline resulted in a penalty in the form of increased licking duration by 200% of the baseline licking during the stimulus presentation. Mice typically learned these rules and completed the pre-training stages in 2-3 days (4-6 sessions of 20 min per animal).

### Training

Each odor discrimination trial lasted 13.2 s, consisting of 3.2 s preloading, 2 s odor presentation/response window (with the last 1 s of preloading as baseline), and a 10 s inter-trial interval, and was initiated by a 200 ms tone before odor onset. For rewarded (S+) trials, water was delivered at the end of the 2 s stimulus if mice either produced double the baseline lick duration during the stimulus/response window or, in the absence of baseline licking, accumulated ≥240 ms of licking distributed over at least 3 of 4 consecutive 500 ms bins. For non-rewarded (S-) trials without baseline licking, correct performance required either no licking or reduced licking (≤80 ms in any one of four bins), and S- errors were not punished so that decision-revision behaviour could be studied. Odor identity (S+ vs S-) was pseudo-randomised within 20-trial blocks to keep rewarded and non-rewarded trials equal, yielding a low probability of repeated identical stimuli over 1200 training trials. Mice performed 150-170 trials per daily session, and motivation was monitored online from lick counts: sessions were terminated when animals showed either persistent, high-rate licking (>40 licks, persistent licking) or markedly low licking (<4-5 licks after reward), ensuring proper motivation levels.

## Statistical analysis

Statistical analyses were performed using custom-written Python scripts, GraphPad Prism 9, and Microsoft Excel. We used one-way and two-way ANOVA, and associated post-hoc tests, student’s t-test, and various non-parametric tests as applicable. The normality of data sets was determined using the Kolmogorov-Smirnov and Shapiro-Wilk tests.

## Acknowledgments

We thank Laboratory of Neural Circuits and Behaviour (LNCB) members and IISER-Pune Biology colleagues for fruitful discussions. We thank staff of *National Facility for Gene Function in Health and Disease* (NFGFHD). Some of the illustrations were created with BioRender.com.

## Author Contributions

N.M.A. supervised all aspects of the project. N.M.A. carried out the study conceptualization and experimental design. S.P. performed optogenetic experiments. S.P., A.S.B., and K.P. performed behavioral experiments. S.P., A.K.N., and S.D. analyzed the data. S.P. and N.M.A. wrote the manuscript.

## Data and Materials Availability

All cumulative data are available in the article figures; further inquiries can be directed to the corresponding author.

